# Identification of a pan-*orthoebolavirus*-reactive antibody from an rVSV-EBOV vaccinated individual

**DOI:** 10.64898/2026.08.28.747769

**Authors:** Paulina Tarnow, Hadas Cohen-Dvashi, Cornelius Rohde, Verena Krähling, Leon Ullrich, Lutz Gieselmann, Hagar Zar Shuker, Shreeya Amatya, Anahita Fathi, Alexandra Kupke, Christoph Kreer, BSL-4 Animal Facility Team, Manuel Koch, Marylyn M. Addo, Stephan Becker, Ron Diskin, Matthias Zehner, Florian Klein

## Abstract

Orthoebolaviruses such as Ebola virus (EBOV), Sudan virus (SUDV) and Bundibugyo virus (BDBV) can cause severe disease with high case-fatality rates. While licensed EBOV vaccines and therapeutic antibodies protect against EBOV infection, no single monoclonal antibody currently provides broad protection across multiple orthoebolaviruses.

Here, we analyzed the humoral immune response of an rVSV-EBOV vaccinee to identify pan-*orthoebolavirus*-neutralizing antibodies. Using BDBV- and SUDV-glycoproteins for single B cell-sorting, we identified B10, which neutralized authentic EBOV and SUDV, with potent activity against SUDV compared with established cross-reactive antibodies. Structural analysis mapped antibody B10 binding to the pan-*orthoebolavirus* conserved GP2-stalk/HR2 region, associated with asymmetric trimer destabilization and spike opening. *In vivo,* B10 showed significant prophylactic efficacy in an EBOV mouse model and partial protection with antiviral activity in a SUDV mouse model.

Together, these findings demonstrate that rVSV-EBOV vaccination induced the development of a broadly orthoebolavirus-neutralizing antibody that holds exeptional therapeutic potential.

## INTRODUCTION

Filoviruses are zoonotic pathogens that can cause severe outbreaks of human disease. Among them, orthoebolaviruses, including Ebola virus (EBOV), Sudan virus (SUDV), and Bundibugyo virus (BDBV), have caused recurrent epidemics with substantial mortality^1^. While EBOV has been responsible for the largest outbreaks to date^2,3^, SUDV and BDBV continue to pose a significant threat to global health preparedness.

EVD can be prevented by licensed vaccines and treated with approved monoclonal antibody (mAb) therapeutics^4–8^, which have substantially improved the control of EBOV disease. However, these countermeasures are largely EBOV-specific, whereas outbreaks caused by other orthoebolaviruses, including SUDV and BDBV, continue to threaten global health security and broadly neutralizing antibodies reactive across multiple orthoebolaviruses remain limited. Thus, the identification of antibodies targeting vulnerable and structurally conserved sites is critical for the development of next-generation therapeutics with broader protective capacity.

Licensed therapeutic mAbs and vaccines are directed against the trimeric surface glycoprotein (GP) of EBOV^9^. After mediating viral attachment and internalization into the host cells, filovirus GP is proteolytically cleaved into the disulfide-linked subunits GP1 and GP2^10–12^. GP1 comprises the heavily glycosylated mucin-like domain (MLD), a glycan cap that shields conserved functional sites and the receptor binding site, whereas GP2 contains the fusion machinery, including the internal fusion loop as well as two consecutive heptad repeat regions (HR1/HR2) and is anchored in the viral membrane via a membrane-proximal external region (MPER) and a transmembrane (TM) domain^13,14^. Importantly, the membrane-anchoring GP2 stalk region (encompassing HR2 and the adjacent MPER), is highly conserved between EBOV, BDBV and SUDV (>90% amino-acid identity) and remains relatively conserved across orthoebolaviruses (about 70% amino-acid identity). This conservation makes the GP2 stalk an attractive target for cross-reactive mAbs and vaccine design. Indeed, stalk-directed antibodies such as BDBV223^15^ demonstrate cross-reactivity, yet breadth across all major orthoebolaviruses remains incomplete.

rVSV-EBOV is a licensed live-attenuated viral vector vaccine expressing EBOV-GP as antigenic target^16^. Although serum neutralization after vaccination is largely EBOV-specific, rare cross-reactive memory B cell clones may nevertheless arise and persist. Studies in other viral systems, including SARS-CoV-2, have demonstrated that ongoing germinal center activity and affinity maturation can progressively expand antibody breadth over time, enabling cross-variant or even cross-sarbecovirus neutralization^17^. Whether rVSV-EBOV vaccination similarly promotes the maturation and persistence of memory B cells targeting conserved structural elements capable of mediating cross-orthoebolavirus neutralization remains unclear.

Here, we analyze PBMCs from an rVSV-EBOV vaccinee to identify vaccine-induced cross-reactive mAbs and to define their breadth and neutralization potency across orthoebolaviruses. We identify B10 as a lead mAb with pan-orthoebolavirus binding and broad neutralizing activity, and map its epitope to the conserved GP2 stalk region. Structural analyses indicate destabilization of the trimeric GP complex as a potential mechanism of neutralization. Finally, we evaluate B10 *in vivo*, demonstrating a favorable pharmacokinetic profile and prophylactic activity in EBOV and SUDV mouse models.

## RESULTS

### Durable cross-reactive memory B cells are detectable nearly two years after rVSV-EBOV vaccination

To characterize the breadth of vaccine-elicited human B cell responses following rVSV-EBOV vaccination, we collected serum and peripheral blood mononuclear cell (PBMC) samples from a healthy individual 21 months after vaccination with 3 × 10^5^ plaque-forming units (PFU) of rVSV-EBOV (single intramuscular injection). For the isolation of GP-reactive memory B cells, we produced recombinant EBOV, BDBV and SUDV-GP and used them as fluorescently labelled baits for single-cell sorting (Fig. 1a). EBOV-reactive viable CD20+IgG+ memory B cells were isolated by single-cell sorting using fluorescein isothiocyanate (FITC)-labeled EBOV-GP bait, yielding a frequency of 0.29%. To isolate BDBV- and SUDV-reactive memory B cells, we used a double staining with allophycocyanin (APC) and FITC-labeled BDBV-GP or SUDV-GP to reduce non-specific binding and false positive sorted events. This resulted in frequencies of 0.33 and 0.091% bait positive IgG+ memory B cells respectively (Fig. S1 and 1b). Thus, cross-reactive memory B cells recognizing heterologous orthoebolavirus GPs were detectable nearly two years after rVSV-EBOV vaccination.

**Fig. 1.**
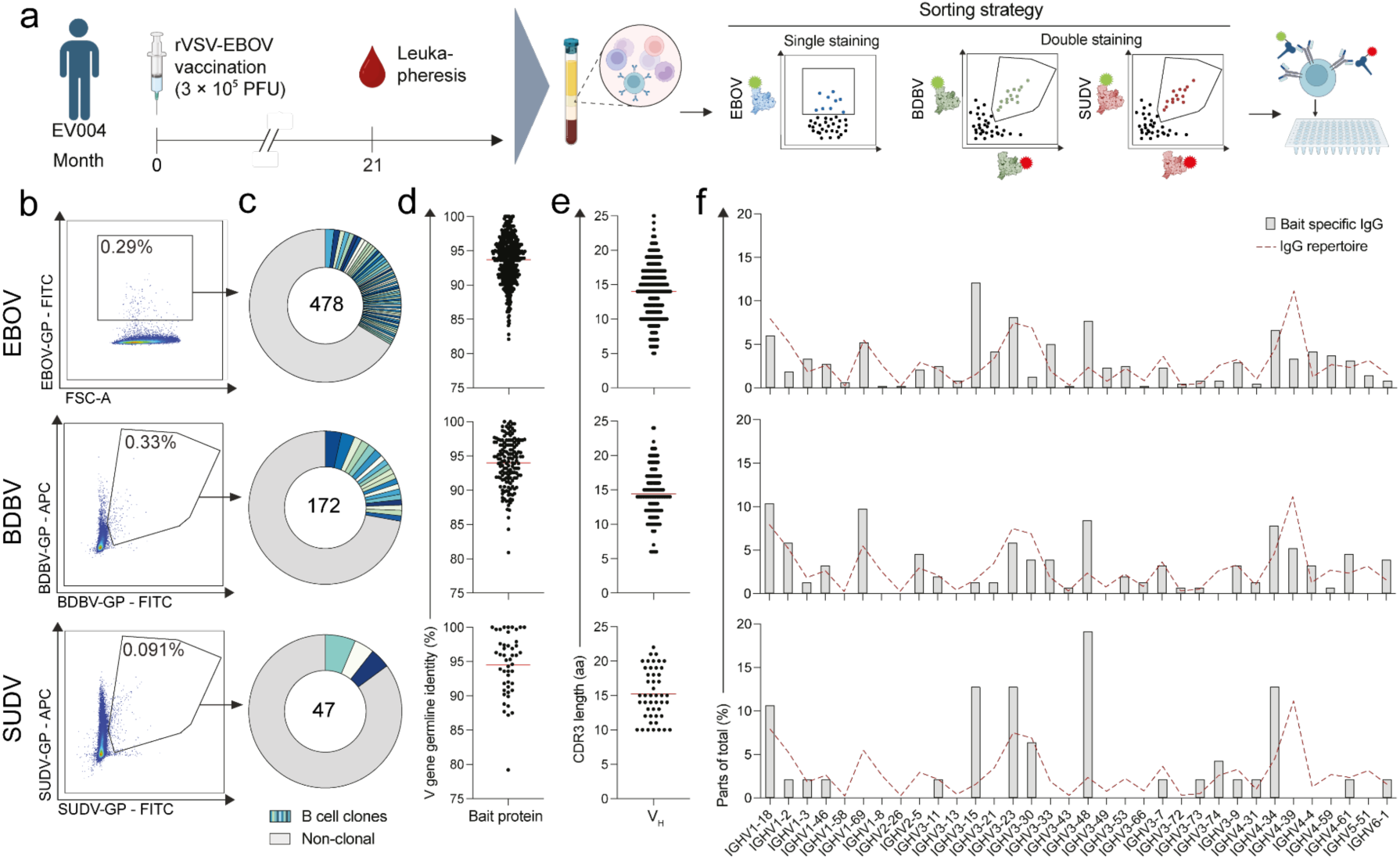
Isolation and sequence analysis of EBOV-, BDBV- and SUDV-GP-reactive memory B cells after rVSV-EBOV vaccination. **a** Sample collection (leukapheresis at month 21 post rVSV-EBOV vaccination) and sorting strategy to isolate EBOV-/BDBV-/SUDV-GP reactive B cells. **b** Single cell isolation of CD19+, CD20+, IgG+ living B cells reactive with fluorescent-labeled EBOV- (top), BDBV- (middle) or SUDV-GP (bottom). Numbers indicate the frequency of GP+ cells within the IgG+ gate. **c** Clonality of isolated B cells, defined by identical VH gene segments usage and CDRH3 region identity greater than 75% amino acids. Distinct blue and green shades represent individual clones, numbers shown within the pie charts indicate the total number of successfully isolated B cell receptors. **d-e** V gene germline identity **(d)** and CDRH3 length **(e)** of isolated B cell receptor sequences, red line indicates mean with standard deviation. **f** Frequencies of VH gene segments of isolated bait specific B cell receptors. Dotted red line indicates the total IgG+ B cell repertoire of the same donor^19^.

For in-depth single B cell and Ig sequence analysis, nested PCR protocols using highly effective primer sets^18^ were applied. After quality checks, we obtained 697 Ig heavy-chain variable (VH) region sequences (EBOV n = 478; BDBV n = 172; SUDV n = 47) and 198 light-chain variable (Vκ/λ) regions of individual B cells. Analysis of VH regions revealed 61 (EBOV), 19 (BDBV) and 3 (SUDV) B cell clones (Fig. 1c; clones defined by identical IGHV gene usage and >75% amino-acid identity in complementarity-determining region heavy chain 3 (CDRH3)). This highlights a polyclonal B cell response across all three glycoproteins, with lower frequency and limited but detectable clonal expansion among SUDV-reactive cells.

The mean V gene germline identity was comparable between different baits and ranged from 93.7% (EBOV) to 94.5% (SUDV) for heavy chains (Fig. 1d). The average CDR3 length in amino acids (aa) of IgH was slightly lower in EBOV sorted B cells (13.9 aa) compared to BDBV (14.4 aa) and SUDV sorted cells (15.2 aa) (Fig. 1e), indicating similar B cell maturation across the bait-specific populations. Despite largely EBOV-specific serum neutralizing activity^19^, these findings indicate that rare cross-reactive memory B cell lineages persist long-term after vaccination.

Next we compared the IgG segments of the EBOV-, BDBV- and SUDV-GP-specific response to the overall B cell repertoire of the same donor using data of unbiased next generation sequencing (NGS) analyses on heavy chains of the memory B cell compartment of the vaccinee^19^. Consistent with our previous analysis, IGHV3-15 was enriched in the EBOV-GP-reactive repertoire (12.0% versus 1.6% in bulk IgG)^19^. Interestingly, IGHV3-15 was also enriched among SUDV-GP-reactive sequences (12.8%), but not for BDBV bait sorted cells (Fig. 1f). A similar enrichment can be seen with IGHV3-48 showing 7.7%, 8.4% and 20% for EBOV, BDBV and SUDV sorted cells respectively compared to 2.4% for the IgG repertoire. In contrast, the frequency of IGHV4-39 was lower in EBOV (3.3%) and BDBV (5.2%) compared to IgG repertoire (11.2%) (Fig.1f). Together, these data reveal selective enrichment of specific VH gene segments within GP-reactive memory B cells, indicating preferential recruitment of defined germline lineages in cross-reactive responses.

Collectively, these findings demonstrate that rVSV-EBOV vaccination can establish a durable memory B cell repertoire that includes cross-reactive lineages capable of recognizing heterologous orthoebolavirus GPs and thus may represent early intermediates of broader neutralizing responses.

### Identification of B10 as a broadly cross-neutralizing antibody with potent activity against authentic EBOV and SUDV

Given the presence of cross-reactive memory B cell lineages, we next aimed to determine whether these cells encoded functionally cross-neutralizing antibodies. To functionally evaluate the cross-reactive memory B cell repertoire, we selected VH/VL pairs from BDBV- and SUDV-GP-sorted memory B cell clones, complemented by a limited number of non-clonal sequences chosen based on somatic hypermutation and CDRH3 length for antibody generation. In total, we cloned and expressed 37 mAbs (Fig. 2a). Of these, 33 mAbs (89%) bound EBOV-GP in ELISA, with half-maximal effective concentration (EC_50_) values ranging from 0.0018 to 24 µg/ml, whereas four mAbs showed no detectable binding. Notably, 18 mAbs displayed cross-binding to other orthoebolavirus GPs (EBOV, BDBV and SUDV) (Fig. 2a). We next analyzed the 33 EBOV-GP-binding antibodies in a lentiviral pseudovirus neutralization assay. Twenty-two mAbs neutralized only a single pseudovirus (EBOV, BDBV or SUDV), with half-maximal inhibitory concentration (IC_50_) values between 6.1 and 900 µg/ml. Six antibodies neutralized two of the three pseudoviruses, and two antibodies (B7 and B10) exhibited cross-neutralization of EBOV, BDBV and SUDV pseudoviruses (Fig. 2b, c). Based on its overall potency and breadth in the pseudovirus panel, B10 was prioritized for follow-up characterization.

**Fig. 2.**
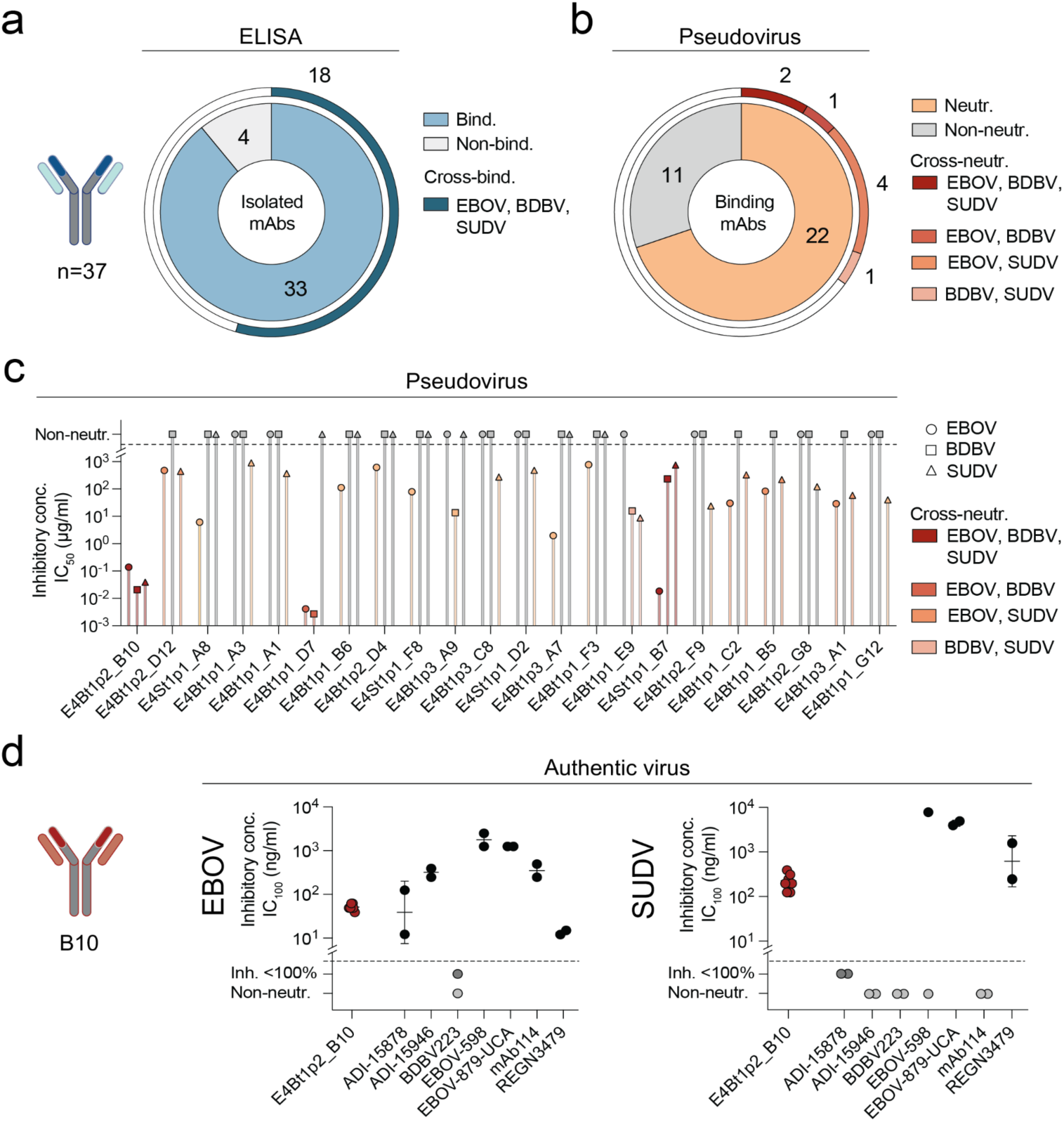
Identification of broadly neutralizing orthoebolavirus mAbs. **a** Recombinant mAbs (n=37) were produced and tested for binding to EBOV, BDBV and SUDV-GP in ELISA. Pie chart indicates binding activity (blue) vs. non-binder (grey); outer ring indicates cross-binding to more than one of the three GPs. Numbers denote the amount of mAbs in each category. **b** The 33 ELISA binding mAbs were tested for neutralization of EBOV, BDBV and SUDV pseudoviruses. Pie chart indicates neutralizing (orange) vs non neutralizing (grey) mAbs, outer ring indicates cross neutralization across two or three pseudoviruses. Numbers indicate antibodies with this neutralization capacity. **c** PSV neutralization potency of individual mAbs shown as half-maximal inhibitory concentration IC_50_ (µg/ml) against EBOV, BDBV and SUDV (symbols). Bars are coloured according to cross-neutralization profile. The dashed line and grey symbols indicate the non-neutralizing mAbs. **d** Neutralization of authentic EBOV (left) and SUDV (right) by the lead candidate B10 (red dots) compared with reference mAbs (black dots). IC_100_ values (ng/ml) are shown; each dot represents an independent replicate measurement for the indicated mAb. Grey symbols denote antibodies that did not reach complete inhibition or were non-neutralizing under the assay conditions.

To determine whether pseudovirus cross-neutralization translated into activity against authentic virus, we performed neutralization assays with authentic EBOV and SUDV. B10 neutralized authentic EBOV and SUDV with mean IC_100_ values of 50.7 ng/ml and 210.3 ng/ml, respectively (Fig. 2d). In parallel, a panel of reference antibodies^20–24^ showed IC_100_ values ranging from 13.4 to 1,767.7 ng/ml against EBOV (with REGN3479^24^ presenting the most potent in this set) and from 623.9 to 7,810 ng/ml against SUDV. Overall, B10 combined broad cross-neutralization with robust activity against authentic virus, including superior potency against SUDV compared with the established cross-reactive mAb REGN3479 (Fig. 2d).

Together, these data establish B10 as a broadly cross-neutralizing antibody with potent activity against authentic EBOV and SUDV.

### Structural analysis reveals B10 binding to the conserved GP2 stalk of EBOV-GP

To map the epitope of B10, we performed antibody competition assays with structurally characterized reference antibodies and found that B10 competes with BDBV223^15^, indicating an overlapping site within the GP stalk region (Fig. S2)

To gain structural insights into B10 recognition of EBOV-GP, we determined a structure of B10 in complex with the EBOV-GP using single-particle cryo-EM (Fig. S3). Three Fab copies were observed bound to a short helical bundle (Fig. S3) within the membrane-proximal GP2 stalk region (Fig. 3a). Interestingly, B10 exclusively utilizes its heavy chain, with no direct contacts contributed by the light chain (Fig. 3b). One helix from the GP’s stalk, hereafter referred to as ‘h1’, is accommodated within a groove formed between CDRH3 and CDRH1/CDRH2 (Fig. 3c), constituting the main binding surface for B10. Val104, a hydrophobic residue at the tip of CDRH3, serves as a central anchoring element and is intercalated between the h1 helix and the adjacent helix ‘h2’ (counterclockwise, along the GP’s axis going from the ectodomain toward the membrane) (Fig. 3c).

**Fig. 3.**
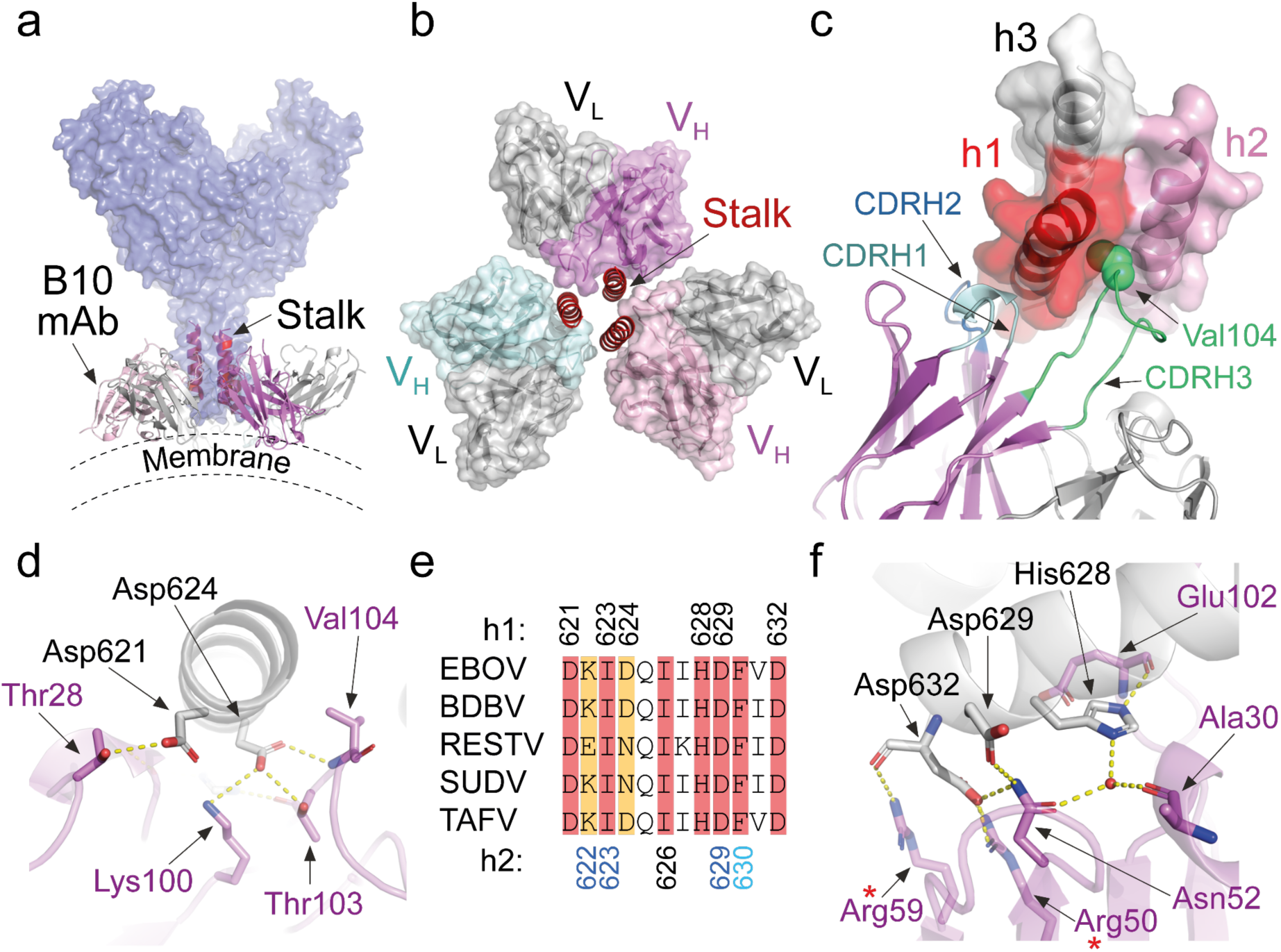
Epitope recognition by B10 mAb. **a** B10 mAb targets the stalk/HR2 region of the GP. The B10 mAb/EBOV complex, illustrated as ribbons, is superimposed on a structure of the EBOV ectodomain (PDB: 6QD8), illustrated as a semitransparent surface. The estimated location of the membrane is schematically illustrated. **b** B10 mAb binds to the GP through its VH domain. The three copies of the B10 VH/VL are shown as semitransparent surfaces in grey (VL) and various colors (VH). The three helices of the spike’s stalk region are shown as red ribbons. **c** One helix from the stalk is cradled in a groove formed between CDRH3 (green) and CDRH1/CDRH2 (shades of blue). Val104 is shown as green spheres. The three helices of the GP’s stalk are annotated as h1, h2, and h3, according to their position with respect to the B10 Fab. **d** A focused view showing how B10 (magenta) recognizes the N’ part of its main h1-epitope (grey). Key residues are noted. Polar interactions are highlighted with yellow dashed lines. **e** The epitope of B10 mAb and its conservation. The stalk/HR2 regions of EBOV, BDBV, RESTV, SUDV and TAFV (Uniprot codes Q05320, B8XCN0, Q89853, Q66814 and Q66810 respectively) that constitute the epitope of B10 mAb are shown. Conserved and semi-conserved (red and orange, respectively) residues that B10 targets are highlighted. Residue numbers that contribute to the epitopes at the h1 or h2 positions are listed at the top and bottom, respectively. At the h2 position, residue numbers that contribute to all Fabs, two Fabs, or to a single Fab are colored black, blue, or light blue, respectively. **f** A focused view showing how B10 (magenta) recognizes the C’ part of its main h1-epitope (grey). Red asterisks mark residues that result from affinity maturation. A water molecule is shown as a red sphere.

Except for Val104 which inserts into a hydrophobic pocket between h1 and h2, the majority of B10 - GP contacts are mediated by polar and charged interactions. These include Asp621, which forms a hydrogen bond with Thr28 (Fig. 3d), and Asp624 that forms a salt-bridge with Lys100 in addition to hydrogen bonds with Thr103 and the main-chain amine of Val104 (Fig. 3d). Interestingly, while Asp621 is conserved among ebolaviruses, residue 624 is an asparagine in SUDV and RESTV (Fig. 3e); however, this substitution is expected to preserve geometry and local interaction through polar contacts.

Further toward the C’, His628 and Asp629, which are both conserved residues (Fig. 3e), engage with either direct or water-mediated polar interactions with B10 (Fig. 3f). The last residue that B10 targets is the conserved Asp632 that forms a salt-bridge with Arg50 and is further engaged by Arg59 that forms a polar interaction with the main-chain carbonyl of Asp632 (Fig. 3e & 3f). Of note, both Arg50 and Arg59 arise from somatic hypermutation, indicating affinity maturation of key contact residues.

Together, these structural data define the conserved GP2 stalk as the epitope of B10 and reveal key contact residues mediating its interaction.

### B10 binding is associated with asymmetric GP stalk rearrangement and head opening

Initial 3D reconstruction attempts of the B10/GP EM map revealed a lack of threefold symmetry (C3); therefore, the map was generated without rotational symmetry being imposed (Fig. S3). This lack of symmetry is evident in the model when the distances between the three copies of B10 are plotted (Fig. 4a), implying that the chemical environment of each binding site is somewhat different. Separating the three B10 Fabs along with their h1 & h2 contact sites into three different models, and then superimposing the three VH/VL pairs reveaed an identical recognition mode of the h1 helices, since the three h1 helices are superimposed perfectly on top of each other (Fig. 4b). The h2 helices, on the other hand, are vertically shifted, and slightly rotated with respect to each other (Fig. 4b). Due to these respective shifts, Lys622, for example, forms polar interactions with CDRH3 of only two B10 Fabs (Fig. 3e & 4b). Also, the hydrophobic pocket that accommodates Val104 (Fig. 3c), is identical on the h1 side in all three binding sites, but is supported by different hydrophobic residues on the h2 side (Fig. 4c). Together, these structural observations are consistent with local destabilization of the GP2 stalk upon B10 engagement.

**Fig. 4.**
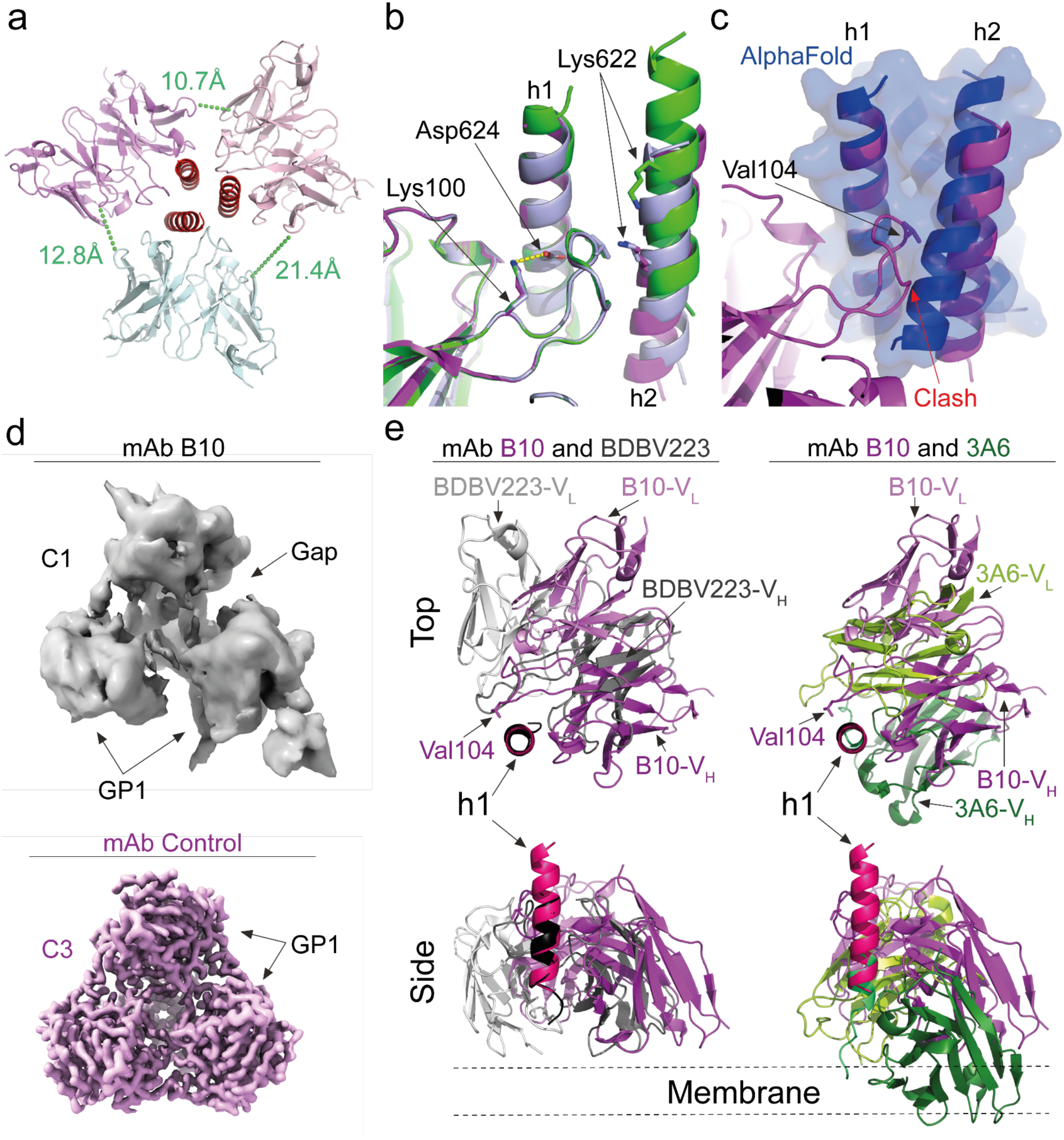
Asymmetric binding of B10 to the GP stalk. **a** The C-C distances between heavy-chain Ser75 and light-chain Ser53. The three copies of the VH/VL pairs are indicated in cyan, pink, and violet. The three helices of the GP’s stalk region are shown as red ribbons. **b** Multiple binding modes of the h2 helix. Three copies of B10 with their h1 and h2 helices (magenta, green, and light purple for the three copies) are shown superimposed on each other, based on the VH/VL pairs. Asp624 of the h1 helix is noted, highlighting perfect alignment of the h1 helices. Lys622 of h2 helices is noted, illustrating their translations with respect to each other. **c** Superimposition of B10 (magenta) on the predicted (AlphaFold3), symmetric stalk of the GP (blue ribbon and semitransparent surface), calculated based on h1 only. A putative clash in this configuration between CDRH3 and h2 is noted. **d** The head region of the GP. EM maps of the head regions, reconstructed without imposing rotational symmetry (C1) from the B10 dataset (top, grey, map level=0.178), and reconstructed with three-fold symmetry averaging (C3) from a dataset collected using the same EBOV-GP batch in the presence of a control Fab (bottom, pink, map level=0.154, deepEMhanced sharpened map). The GP1 domains are annotated, as well as a gap between the domains. **e** Previously identified anti-stalk antibodies. Structures of 3A6 (PDB: 7RPU) and BDBV223 (PDB: 6N7J) Fabs in complex with h1 peptide are shown for comparison with mAb B10. The structures were superimposed based on the h1 helices to illustrate how this region is engaged by the different antibodies. The top panel shows ‘top’ views of the antibodies, which are represented as ribbons colored in shades of magenta, grey, and green for B10, BDBV223, and 3A6, respectively. Val104 in the CDRH3 of B10 is indicated for orientation. The lower panel shows ‘side’ views of the structures, with the estimated location of the membrane schematically indicated.

Fitting B10 onto an idealized threefold-symmetric stalk model generated by AlphaFold3 suggests that conformational adjustments of the coiled-coil stalk would be required to accommodate B10 binding (Fig. 4c). First, the helices must move outward, away from each other, to allow the intercalation of Val104 into the otherwise compact core of the coil (Fig. 4c). Moreover, due to a clash with CDRH3, the h2 helices rotate such that they align more parallel to each other, losing the natural twist of the coiled coil (Fig. 4c). Combined, these two movement components likely destabilize the interaction of the helices in the stalk. We speculate that under these conditions, the available energetic minima for packing the three helices require one of them to slide with respect to the others (Fig. 4b), resulting in the observed asymmetry. Regardless of the exact mechanism, this asymmetry has a profound effect on other regions of the GP.

The EM map of the three Fabs did not include density for the GP1 domains that comprise the head region of the GP, indicating flexibility between the head and the stalk regions. To reveal the structure of the head, we performed particle picking using templates generated from a known structure of the EBOV-GP (PDB: 6QD8) and reconstructed a map for the head region (Fig. S4). Despite having a substantial amount of data and testing various classification strategies, we were only able to generate a low-resolution, non-symmetric map for the head region (Fig. S4), indicating structural heterogeneity. Although low resolution, the map clearly reveals relative motion and opening of the GP1 domains (Fig. 4d). Moreover, 3D classes imply the existence of potentially other wide-open states of the GP.

This opening of the GP is associated with the binding of B10, since single-particle cryo-EM analysis of the exact same protein batch of EBOV-GP in the presence of a non-binding Fab (Fig. S5) readily provides a high-resolution, three-fold symmetric map for the head region (Fig. 4d). Collectively, these findings indicate that B10 engagement is accompanied by asymmetric stalk rearrangement and increased head flexibility, consistent with a destabilizing mode of neutralization.

Interestingly, structural superposition with the stalk-directed antibodies BDBV223 and 3A6^15,25^ highlights that, despite targeting the same general stalk/HR2 region, B10 engages the h1 helix at a distinct angle centered around Val104, highlighting a divergent molecular strategy for targeting this conserved epitope (Fig. 4e).

Collectively, these structural data support a model in which B10 binding destabilizes the GP stalk and promotes asymmetric head opening.

### *In vivo* pharmacokinetics and prophylactic efficacy of B10 in EBOV and SUDV mouse models

To evaluate the *in vivo* antiviral activity of B10, we first analyzed its pharmacokinetic properties in NRG mice. Following intravenous administration of 0.5 mg B10 to mice, serum IgG levels declined gradually over 24 days and were comparable to the kinetics of the HIV reference antibody 10-1074^26^ and 3BNC117^27^ (Fig. 5a), indicating favorable half-life properties over the observation period.

**Fig. 5.**
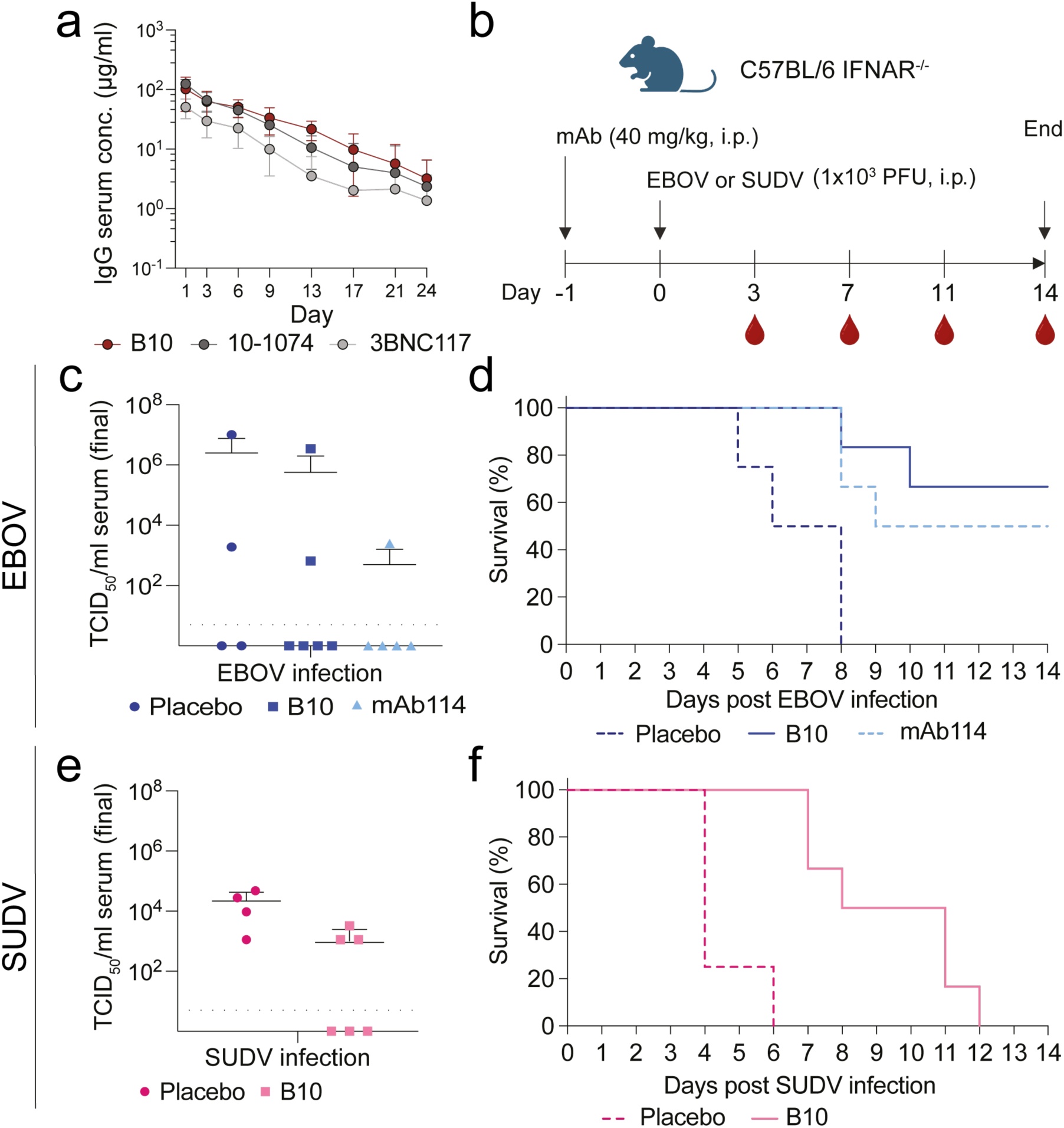
Pharmacokinetics and *in vivo* efficacy of B10 in EBOV and SUDV challenge models. **a** Serum human IgG concentrations over time following a single intravenous administration of purified wild-type antibodies (B10, 10-1074, and 3BNC117; 0.5 mg in PBS; n = 4) in human FcRn-transgenic mice, quantified by human IgG capture ELISA**. b** Experimental timeline for challenge experiments in C57BL/6 IFNAR^−/−^ mice (n = 6; placebo n = 4).Mice received a single intraperitoneal dose of the indicated mAb (40 mg/kg) on day −1 and were infected intraperitoneally with 1×10^3^ PFU of EBOV or SUDV on day 0. Blood was collected on days 3, 7, 11 and 14 or at humane endpoint. **c** Serum viral titers (TCID₅₀/ml) in EBOV-infected mice at humane endpoint or study termination (d14) following treatment with placebo, B10, or mAb114**. d** Survival of EBOV-infected mice following prophylactic antibody administration. **e-f** As in (**c**) and (**d**), but with SUDV-infected mice.

Next we investigated the prophylactic efficacy of B10 in IFNAR^−/−^ mice that lack type I interferon receptors and thereby are highly susceptible to filovirus infections^28^. C57BL/6 IFNAR^−/−^ mice received i.p. a single prophylactic dose of B10, mAb114 or placebo (DZIF10c) (40 mg/kg) one day prior to i.p. infection with 1×10^3^ PFU EBOV or SUDV (Fig. 5b). To reflect the severity of disease progression a clinical score was determined daily. In EBOV infected mice, the terminal clinical score was significantly lower on average in B10 (average score: 5.0) or with the licensed monoclonal anti-EBOV Ab mAb114^22^ (average score: 8.3) treated animals compared to placebo (average score: 14.3), whereby B10 increased survival to 66.7%, exceeding the survival observed with mAb114 (50%) (Fig. S7a and 5d). In line with this protection, body weight and temperature remained stable in B10- and mAb114-treated animals. In contrast, mice that reached humane endpoint showed pronounced weight loss (Fig. S7b) and developed terminal hypothermia (Fig. S7c). Post mortem analysis of serum, spleen and liver samples revealed reduced EBOV titers (Fig. 5c and S7f). Moreover, consistent with improved survival, B10 treatment reduced EBOV RNA levels in spleen and liver (Fig. S7f, filled symbols compared to unfilled symbols).

After SUDV infection, B10 showed measurable *in vivo* activity in this stringent model, delaying disease progression and extending time to humane endpoint (Fig 5f). Despite the absence of full protection, B10-treated animals exhibited lower terminal clinical scores (average score B10: 13.8; placebo: 16.0) and a prolonged time to endpoint (up to 12 days versus 6 days in the placebo group; Fig. 5d, Fig. S7a). Notably, B10 also delayed the onset of weight loss compared with placebo (day 4 versus day 2) (Fig. S7d). In-line with that, B10-treatment led to reduced titers of infectious SUDV in final serum samples (TCID_50_ 0.9×10^3^/ml versus 2.1×10^4^/ml, mean values) (Fig. 5e) and SUDV RNA levels in spleen and liver (Fig. S7g).

Together, these findings demonstrate that B10 confers robust prophylactic protection against EBOV and measurable *in vivo* efficacy against SUDV.

## DISCUSSION

Licensed antibody therapeutics such as Ansuvimab^TM^ (mAb114) and the antibody cocktail Inmazeb^TM^ (REGN-EB3) are effective treatments for EVD, but do not provide efficient protection against other clinically relevant orthoebolaviruses, including SUDV. While additional vaccines against other orthoebolaviruses are in development, broadly reactive therapeutics that could be applied against distinct orthoebolavirus outbreaks are not yet available. Therefore, there remains a medical need for cross-reactive mAb therapeutics with broad activity against EBOV, BDBV and SUDV to improve outbreak preparedness.

Although rVSV-EBOV vaccination can induce a diverse antibody repertoire with detectable cross-reactive binding, broad neutralization was restricted to a very limited subset of antibodies^19^. Here, we isolated a potent neutralizing antibody (B10) from a rVSV-EBOV vaccinated individual, showing potent neutralization activities against authentic EBOV and SUDV *in vitro*. In a prophylactic *in vivo* model, B10 administration provided protection comparable to mAb114 against EBOV challenge, and exceeded the survival observed with mAb114 in this experimental setting, supporting the *in vivo* relevance of B10-mediated neutralization. Importantly, even clinically validated antibodies such as mAb114 do not achieve complete protection in these experimental settings, underscoring the relevance of identifying additional antibodies with complementary breadth and mechanisms of action.

Our findings indicate that rVSV-EBOV vaccination can induce rare cross-reactive memory B cell lineages that may not be apparent at the serum level but are capable of giving rise to broadly neutralizing antibodies. This observation aligns with the idea that conserved structural elements can be targeted by a minor fraction of the memory repertoire, even when serum neutralization remains largely strain-specific^29^.

The identification of B10 as a GP2 stalk-directed antibody places our findings within the broader context of stem-targeting antibodies. Multiple studies have highlighted the GP_1,2_ stalk region as a promising target for cross-reactive vaccine and mAb design, given its high sequence conservation and functional constraint between different orthoebolaviruses. Notably, King et al. BDBV223 established the GP2 stalk as a cross-reactive site of vulnerability and provided a conceptual framework for stem-targeted antibody design. Recent work on the stalk-MPER-directed antibody 3A6 further underscores the therapeutic potential of this region by providing structural evidence for epitope access through GP “lifting” from the virion membrane and demonstrating strong *in vivo* efficacy against EBOV at low doses levels^25^. Importantly, however, stalk-MPER-directed mAbs can display restricted breadth: 3A6 does only neutralize EBOV, and the stalk-related antibody BDBV223, despite targeting this conserved region, shows activity limited to EBOV and BDBV^15,25^. Our data extend these observations by demonstrating that distinct binding geometry within the same conserved stalk region can expand neutralization breadth to include SUDV.

Together, these findings highlight that high sequence conservation alone does not necessarily translate into breadth and that differences in epitope accessibility and recognition geometry can critically influence cross-neutralization. Structural comparison suggests that B10 engages the GP2 stalk with a distinct angle and an additional loop structure relative to these antibodies enabling the cross-reactivity with SUDV. These observations support the notion that subtle differences in binding orientation and contact topology within a conserved epitope can translate into meaningful differences in cross-neutralization breadth.

In contrast to models proposed for BDBV223 and 3A6 that involve pronounced stalk displacement or stabilization of a lifted GP conformation, B10 binding appears compatible with the prefusion architecture without requiring large-scale bending or lifting. We therefore propose that B10 may neutralize EBOV as well as SUDV by destabilizing the GP spike or restricting the conformational transitions required for productive receptor engagement and membrane fusion. This mechanism differs from previously described stalk-directed antibodies and suggests that multiple structural solutions exist for targeting the GP2 stem. Although the precise sequence of conformational events remains to be defined, B10-induced GP destabilization and opening may expose cryptic vulnerability sites of the GP that are otherwise concealed. Therefore, B10 may potentiate humoral responses in vivo and synergize with specific mAbs targeting such cryptic sites.

*In vivo*, B10 did not achieve complete protection in mice against SUDV infection in this stringent IFNAR^−/−^ model; however, B10 application consistently delayed disease progression and reduced clinical burden. Partial protection in such highly susceptible settings may still reflect biologically meaningful antiviral activity, particularly in the context of antibodies targeting conserved but functionally constrained epitopes.

While our study provides structural and functional evidence for cross-orthoebolavirus neutralization by B10, several aspects warrant further investigation. The antibody was isolated from a single vaccinee, and its in vivo activity was assessed in mouse models, which may not fully recapitulate human disease. Moreover, although B10 demonstrated robust prophylactic efficacy against EBOV, protection against SUDV remained partial in this stringent setting. Future studies in additional donors, advanced animal models and vaccine settings will be important to define the therapeutic potential and translational relevance of GP2 stalk-directed antibodies.

Taken together, our data support the GP2 stalk region as an important site of vulnerability and suggest that modest differences in binding orientation and mechanism can translate into meaningful differences in cross-neutralization breadth of orthoebolaviruses. These findings provide a framework for designing next-generation antibodies and immunogens that preserve the potency of stalk-targeting therapeutics while extending activity across multiple pathogenic orthoebolaviruses.

## RESOURCE AND DATA AVAILABILITY

### Materials availability

Requests for materials will be fulfilled by the corresponding author upon request. Transfer of antibodies, protein constructs, and sample material might require a Material Transfer Agreement (MTA) for non-commercial usage. Human-derived samples are subject to data protection regulations and can only be shared in accordance with applied study protocols and approval by the Ethics Committee of the University of Cologne.

### Data and code availability

Amino acid sequences used in this study were obtained from publicly available datasets. All results generated in this study are presented within the article and its Supplementary Information. No original code was generated for this work. Coordinates and electron microscopy (EM) maps were deposited and are available at the Protein Data Bank (PDB)/Electron Microscopy Data Bank (EMDB) under accession codes 9TGC/EMD-55896. Further details necessary to reproduce or reanalyze the data can be obtained from the lead contact upon request.

## ACKNOWLEDGMENTS

We thank the study participant for taking part in this study and for their time and commitment. We are grateful to all members of the Klein Lab, our collaboration partners, and the animal facilities for their support and valuable discussions. We thank Anna Schmitt, Susanne Salomon, and Tina Bresser for laboratory management and technical assistance. We also thank Jacqueline Knüfer and Ricarda Stumpf as well as the sample processing laboratory for their outstanding support with sample collection, processing, and purification. Elements of Figure 1a, 2a, 2d and 5b were created with BioRender.com. This work was supported by institutional funding from the University Hospital of Cologne.

## AUTHOR CONTRIBUTIONS

Conceptualization, P.T., M.Z., and F.K.; Methodology, P.T., C.R., V.K., M.A., S.B., R.D., M.Z., and F.K.; Investigation, P.T., H.C., C.R., V.K., L.U., L.G., H.S., A.F., C.K., M.K.,; Visualization, P.T., R.D. and M.Z.; Formal analysis, P.T., H.C., C.R., V.K., C.K., M.Z., and F.K.; Writing original draft, P.T., R.D., M.Z., and F.K.; Writing review & editing, all authors; Funding acquisition, R.D., M.Z., and F.K.; Project administration, P.T., M.Z., and F.K.; Supervision, S.B., R.D., M.Z., and F.K.

## ETHICS APPROVAL

All animal experiments were approved by the relevant local authorities (animal welfare committee; Regierungspräsidium Gießen, Germany; approval number: AZ V54-19 c 20 15 h 01 MR 20/7 Nr. G 64/2024) and conducted in accordance with the Federation of European Laboratory Animal Science Associations (FELASA) and Gesellschaft für Versuchstierkunde / Society of Laboratory Animal Science (GV-SOLAS) guidelines, the German Animal Welfare Act, and Directive 2010/63/EU.

Experiments involving infectious Ebola virus (EBOV) and Sudan virus (SUDV) were performed in a registered and ethics accredited biosafety level 4 (BSL-4) facility at Philipps University Marburg, Germany, in accordance with international biosafety regulations and with appropriate regulatory approvals.

Human samples were obtained following written informed consent and with approval from the Institutional Review Board of the University of Cologne, Germany (INA; 16–054), in accordance with Good Clinical Practice guidelines.

## SUPPLEMENTARY INFORMATION

**Fig. S1.**
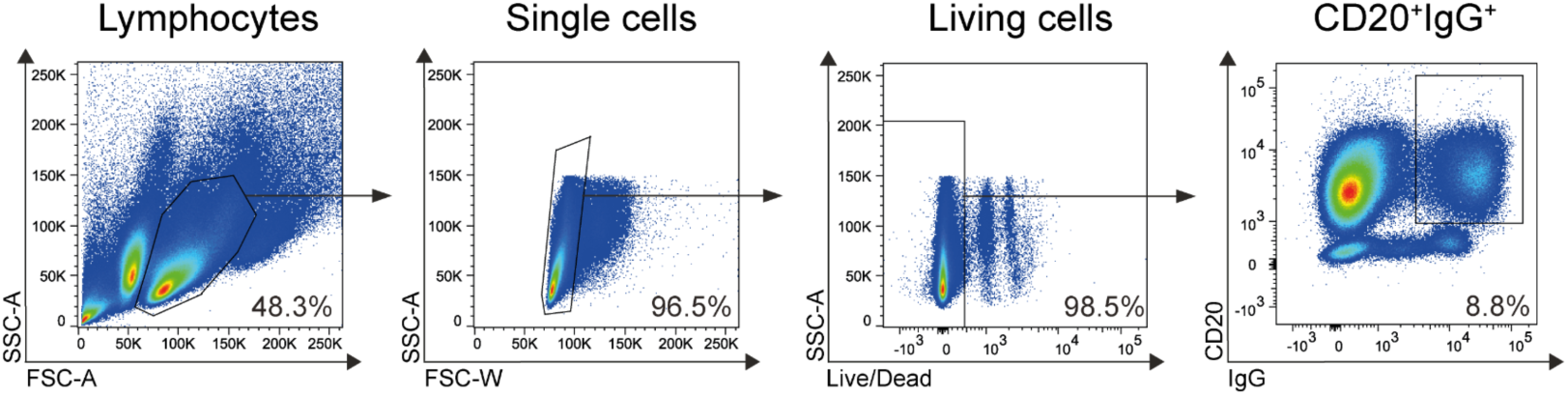
Gating strategy. PBMCs from individual donors were enriched for CD19⁺ B cells by magnetic separation and stained with viability dye, anti-CD20 and anti-IgG, together with fluorescently labeled filovirus GP bait proteins. Representative gating workflow used for single-cell sorting: lymphocytes were identified by FSC-A/SSC-A, doublets were excluded by FSC-W/SSC-A, live cells were selected by Live/Dead staining, and CD20⁺IgG⁺ B cells were gated for subsequent selection of bait-positive cells.

**Fig. S2.**
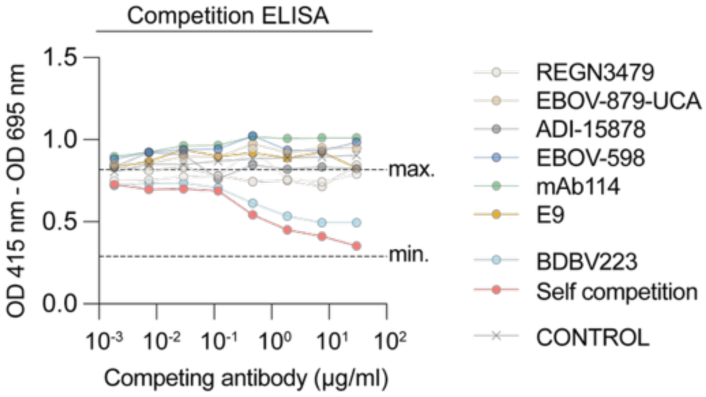
Competition ELISA. EBOV-GP was immobilized on ELISA plates and incubated with serial dilutions of competitor antibody B10 before addition of the indicated test antibody. Binding was normalized to the no-competitor condition (max. dashed line). Self-competition served as a positive inhibition control, and an irrelevant antibody as a negative control. Competition with reference mAbs (REGN3479, EBOV-879, ADI-15758, ADI-15999, mAb114 and E9) was used to assign antibodies to epitope sites based on reduced binding relative to the no-competitor condition.

**Fig. S3.**
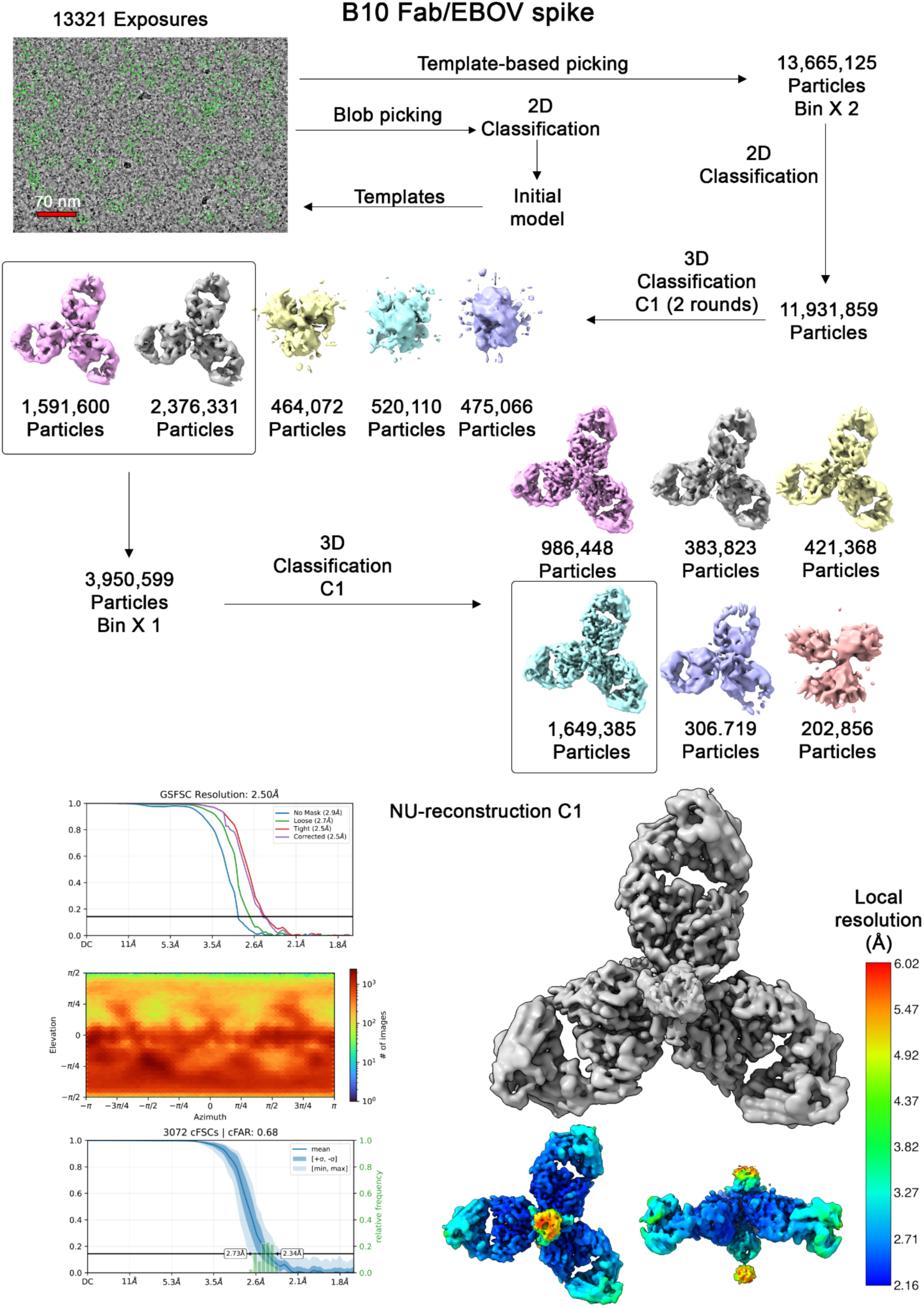
Reconstruction of an EM map for the B10/EBOV complex. The EM map reconstruction pipeline is illustrated. The initial model was generated from blob-picked particles. Template-based picking was used subsequently. No symmetry constraints were imposed for the final reconstruction.

**Fig. S4.**
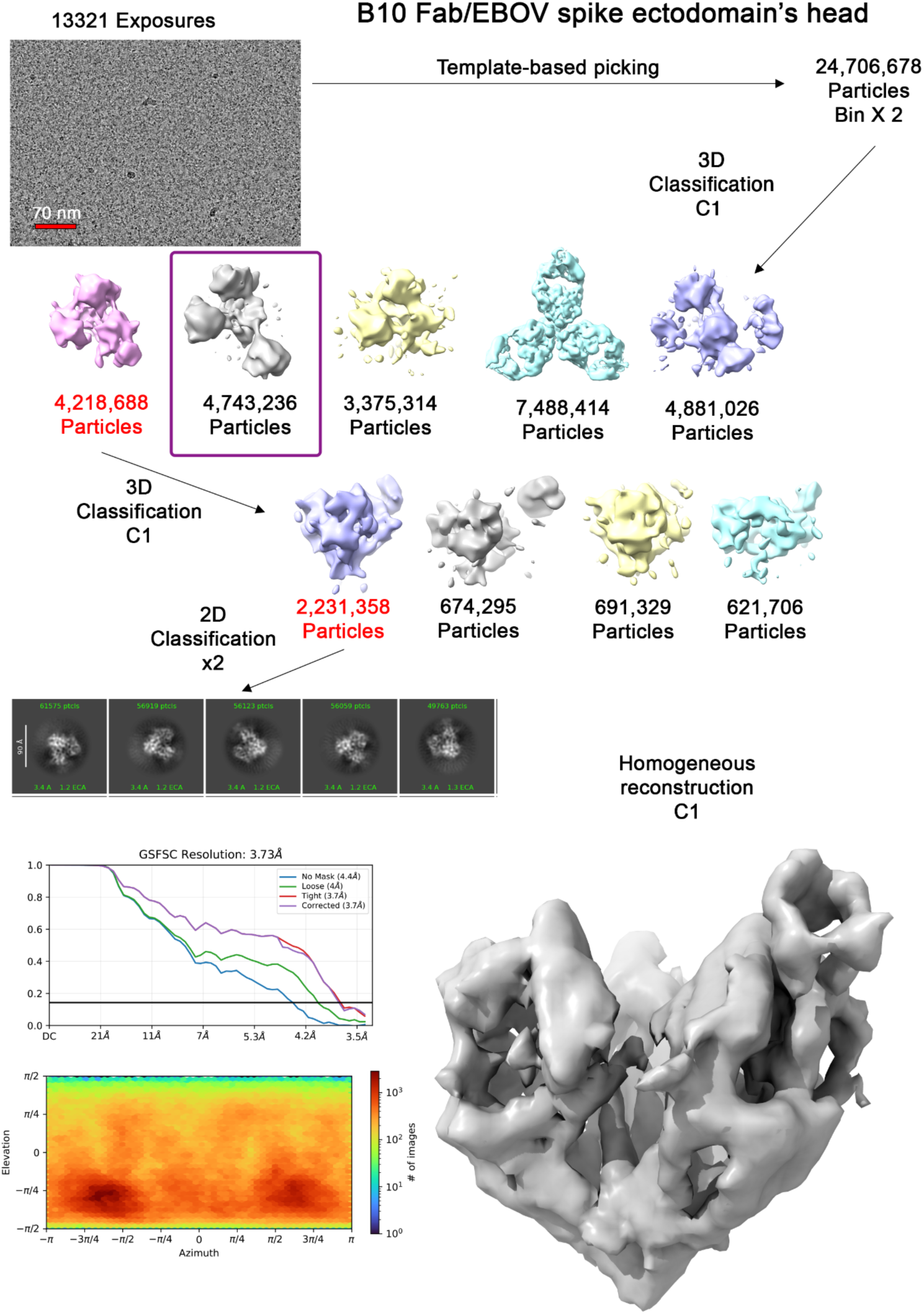
Reconstruction of an EM map for the head region of the B10/EBOV spike complex. This is a representative pipeline out of several attempts that involved various combinations of 2D and 3D classification strategies. Classes with red notations were carried forward. One 3D class in a box shows a wide-open configuration of the spike. The resulting map has a low resolution and overall low quality.

**Fig. S5.**
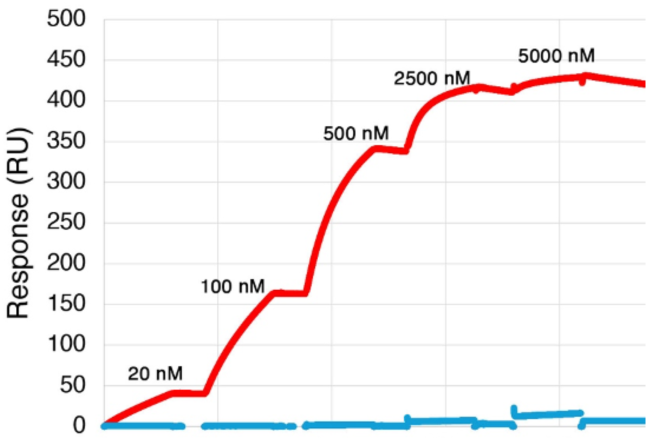
Single-cycle kinetic analysis using surface plasmon resonance (SPR). B10 and E9 (control) Fabs (red and blue curves, respectively) were injected as analytes at the indicated concentration series over covalently immobilized EBOV spike. Response (RU) over time. This assay configuration does not allow avidity.

**Fig. S6.**
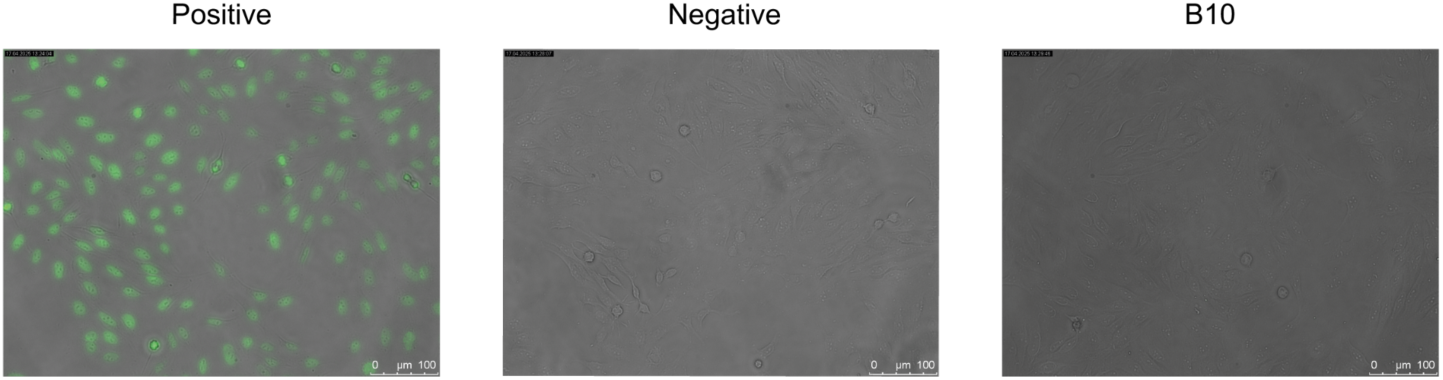
HEp2 Autoreactivity Assay. Reactivity of B10 against HEp-2 cells together with a negative and positive control. The Antibody was tested at a concentration of 100 µg/ml in a single experiment. Data is presented from a single experiment.

**Fig. S7.**
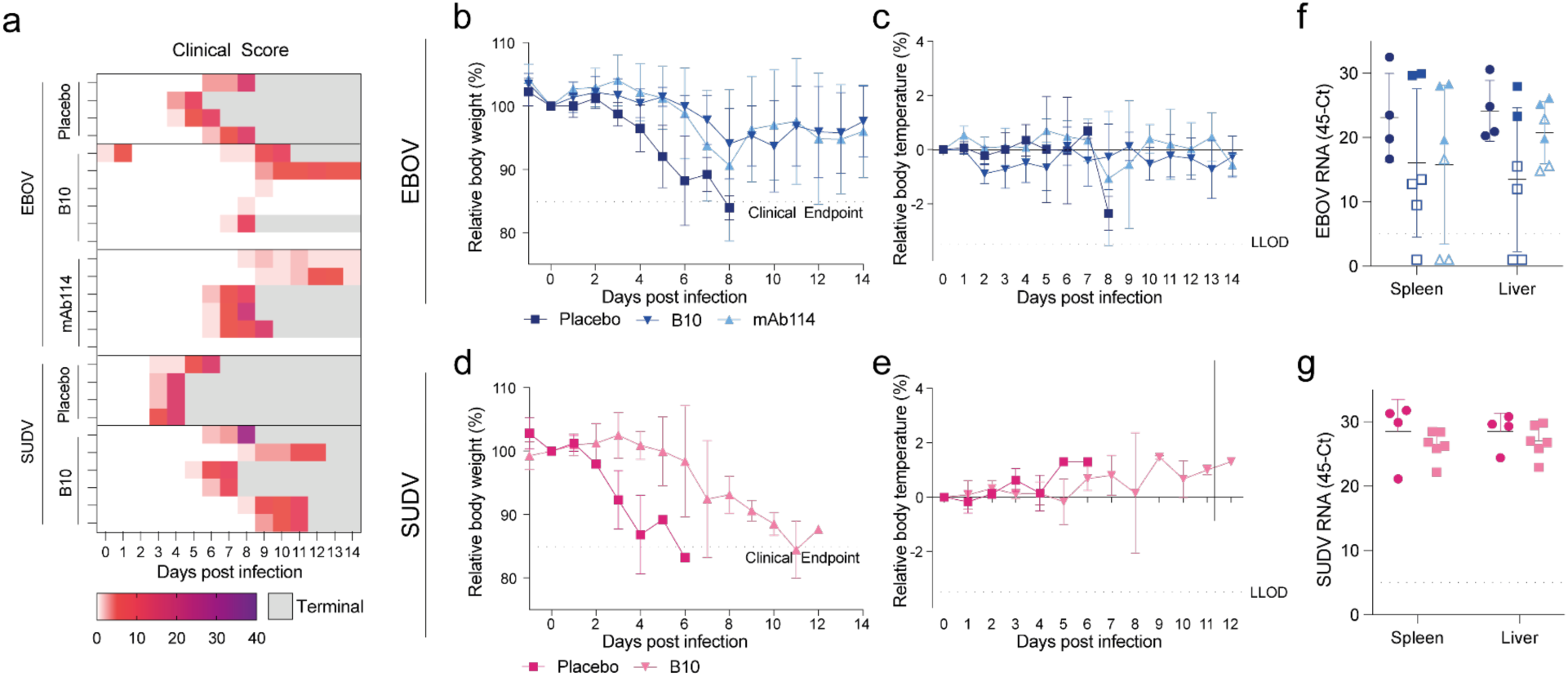
Pharmacokinetics and *in vivo* efficacy of B10 in EBOV and SUDV challenge models. **a** Clinical scores of IFNAR^−/−^ mice following i.p. infection with 1×10^3^ PFU EBOV or SUDV. Mice were treated one day prior to infection (day -1) with 40 mg/kg B10, mAb114 (EBOV control), or DZIF10c (placebo). Heatmaps show cumulative clinical scores. Gray fields indicate animals reaching humane endpoint (terminal). **b** Relative body weight (%) of EBOV-infected mice over 14 days post infection (p.i.). The dashed line indicates clinical endpoint. **c** Relative body temperature of EBOV-infected mice. The dashed line indicates the Lower Limit of Detection (LLOD) **d** Relative body weight (%) of SUDV-infected mice over 14 days p.i (see also **b**) **e** Relative body temperature of SUDV-infected mice (see also **c**) **f** EBOV RNA levels (top) and SUDV RNA levels (bottom) in spleen and liver at day 14 p.i. or at humane endpoint, determined by RT-qPCR. Filled symbols indicate animal survival compared to unfilled symbols. The dashed line indicates the LLOD.

**Table S1.**
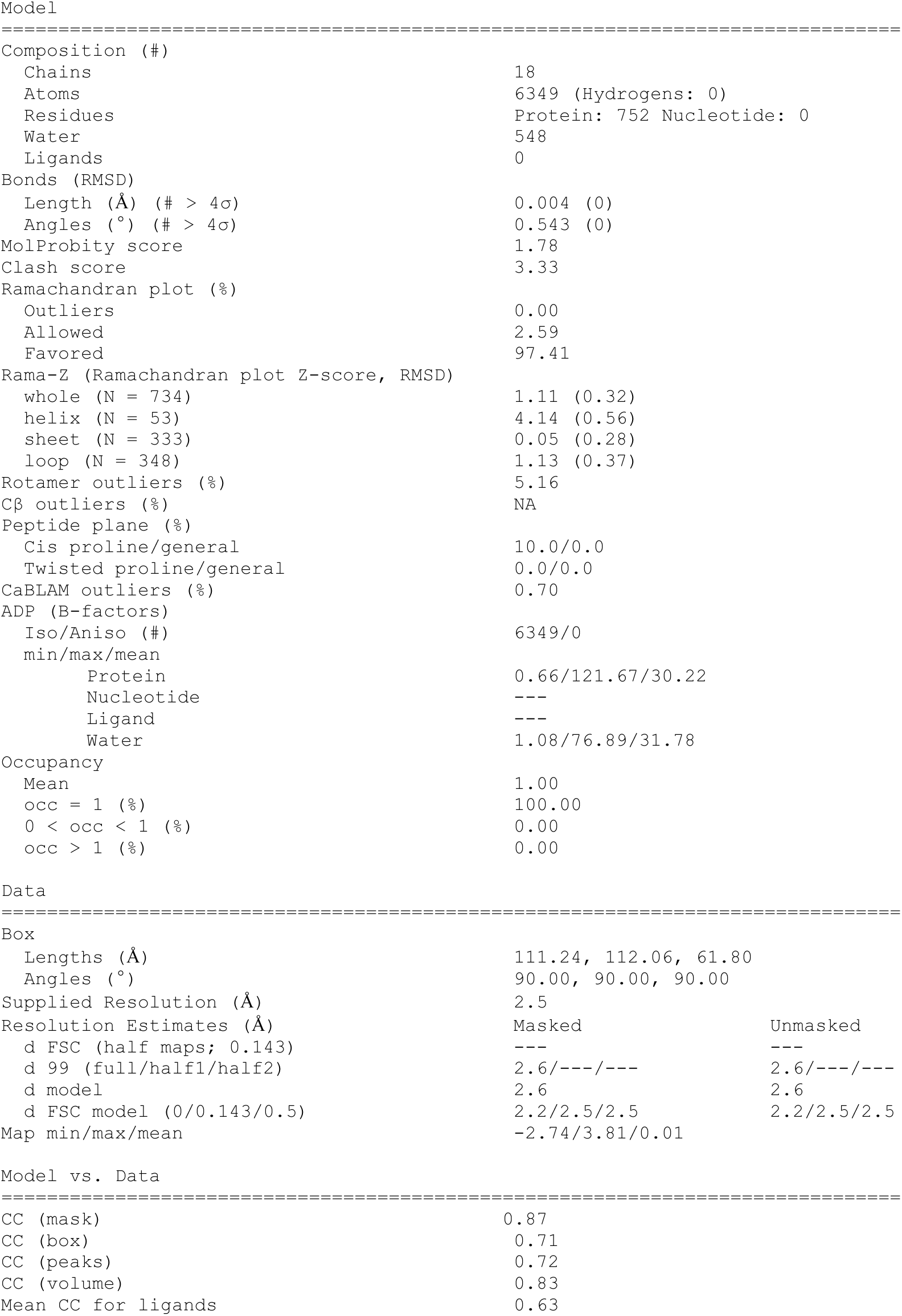
Model refinement statistics for the B10/EBOV complex.

## METHODS

### Study participants and collection of clinical samples

The individual included in this study had previously participated in the Phase I clinical trial evaluating the safety, tolerability, and immunogenicity of the Ebola virus vaccine rVSVΔG-ZEBOV-GP (NCT02283099) and received a dose of 3×10⁵ PFU Serum and leukapheresis samples were obtained from this participant (INA; 16-054, approved by the Institutional Review Board of the University of Cologne, Germany). Peripheral blood mononuclear cells (PBMCs) were isolated by density gradient centrifugation using HistoPaque and cryopreserved in fetal bovine serum (FBS) containing 10% dimethyl sulfoxide (DMSO) (all from Merck, Darmstadt, Germany) at −150 °C. Serum and plasma samples were stored at −80 °C. The participant provided written informed consent prior to study enrollment, and all study procedures were conducted in accordance with Good Clinical Practice guidelines.

### Cell lines and viruses

HEK293T cells (ATCC) were grown in Dulbecco’s Modified Eagle Medium (DMEM) (Thermo Fisher) supplemented with 10% FBS (Sigma-Aldrich), 1 mM sodium pyruvate (Gibco), and 2 mM L-glutamine (Gibco) at 37 °C in a humidified atmosphere containing 5% CO₂. HEK293-6E cells (National Research Council of Canada) were cultured in FreeStyle 293 Expression Medium (Life Technologies) supplemented with 0.2% penicillin/streptomycin, maintained under continuous agitation (90-120 rpm) at 37 °C with 6% CO₂. All cell lines were of female origin and were not additionally authenticated.

For structure generation, HEK293F suspension cells (Invitrogen) were maintained in FreeStyle Medium (Life Technologies) at 37 °C with 8% CO_2_, shaking at 130 rpm in a humidified incubator. Transfections were performed at a concentration of about 1×10^6^/ml, using PEI-Max (Polysciences) at a ratio of 3:1 with 1 µg DNA ml^−1^ cell suspension.

For the efficiency study *in vivo*, Vero C1008 cells (ATCC CRL-1586) were cultured as described elsewhere^30^. SUDV Boniface (GenBank accession number FJ968794.1) and EBOV Mayinga (GenBank: NC_002549) were propagated and titrated in Vero C1008 cells as described elsewhere. All experiments with Filoviruses were carried out in the Biosafety Level (BSL)-4 laboratory of the Marburg University, Germany.

### Isolation of single anti-orthoebolavirus-reactive B cells

B lymphocytes were enriched by magnetic cell separation (MACS) using CD19 microbeads (Miltenyi Biotec) following manufacturer’s instructions. CD19+ B cells were stained with 4’,6-Diamindin-2-phenylindol (DAPI, Miltenyi Biotec), CD20 (BD, clone 2H7, Alexa Fluor 700), IgG (BD, clone G18-145, PE) and either Alexa Fluor 488- and Alexa Fluor 647-labeled EBOV, BDBV or SUDV-GP (10 µg/ml). DAPI−, CD20+, IgG+, and double positive EBOV-, BDBV- or SUDV-GP cells were single cell sorted into 96-well plates using an FACSAria Fusion (BD). Wells contained 4 µl lysis buffer (0.5× phosphate-buffered saline (PBS), 0.5 U μl^−1^ RNasin (Promega), 0.5 U μl^−1^ RNaseOut (Thermo Fisher) and 10 mM DTT (Thermo Fisher)) and were immediately stored at −80 °C after sorting until further processing.

### B cell receptor amplification and sequence analysis

The PCR amplification and sequencing of antibody heavy and light chain genes from sorted single cells was performed as described previously^31^. Briefly cDNA was generated using random hexamer primers (Thermo Fisher) and Superscript IV (Thermo Fisher). Amplification of B cell receptor heavy and light chain sequences was conducted in a semi-nested PCR using human V gene segment specific forward and IgG constant region-specific reverse primers together with Platinum Taq Polymerase (Thermo Fisher)^18^. E

### Cloning and production of mAbs

Variable regions of selected antibodies were ordered as gene fragments without endogenous leader signal from IDT as eBlocks. Sequences were cloned into IgG1, IgK, or IgL expression vectors^35^ using sequence- and ligation-independent cloning (SLIC) with T4 DNA polymerase (New England Biolabs) and transduced into Escherichia coli DH5α as described previously^31,36^. Evaluation of 3-6 colonies by PCR and gel electrophoresis was performed and subjected to Sanger sequencing. Amplification of plasmids containing correct antibody sequences was performed by midi preparation (Macherey Nagel kit) according to the manufacturer’s protocol and stored at 4 °C.

Production of mAbs was performed by co-transfection of heavy and light chain-encoding expression plasmids into HEK293-6E cells at a concentration of 0.8×106 cells/ml with branched PEI (25 kDa, Sigma-Aldrich) as described previously^31,36^. The cells were cultured in FreeStyle 293 medium (Thermo Fisher) with 0.2% penicillin/streptomycin at 37 °C and 6% CO_2_ in a shaking incubator at 110 rpm for 5-7 days.

For structure generation, EBOV ectodomain, mAb B10 and mAb E9 were expressed in HEK293F cells. For mAb expression, vector encoding the heavy and light chains were co-transfected. Media were collected after 6 days of incubation, and IgGs were captured using protein-A affinity chromatography (Cytiva). IgGs were digested using Papain (Sigma-Aldrich) with an enzyme-to-protein ratio of ∼1:20. Digestion proceeded for 60 minutes at 37 °C in a buffer containing 20 mM cysteine-HCl (Sigma-Aldrich) and 10 mM EDTA, titrated to pH 7.0. Fabs were separated from Fc fragments by collecting the flow-through fraction from a protein-A column, followed by size exclusion chromatography (SEC) on Superdex 200 10/300 increased column (Cytiva).

### Vector construction

For structure generation, ectodomain of Ebola virus (EBOV; PDB Seq ID: 5JQB_A) GP was chemically synthesized (Genscript) to eliminate the mucin-like region and to include a C-terminal trimerization domain of T4 fibritin (foldon) followed by a His-tag as previously reported^37^.

### Glycoprotein production and purification

The ectodomains (Sudan virus, WEY06878, aa:33-650 T456P; Bundibugyo virus, YP_003815435, aa: 33-650; Ebola virus, NP_066246, aa: 33-650) were amplified from synthetic gene plasmids using specific PCR primers. The PCR products were digested with the appropriate restriction enzymes (NheI and BglII) and cloned into a modified sleeping beauty transposon expression vector containing a N-terminal BM40 signal peptide sequence and a C-terminal T4 fibritin foldon followed by a Twin-Strep-tag. HEK293 cells were co-transfected with the transposase (0.2 g) and the different ectodomain containing plasmids (1.8 g). The transfected HEK293 were selected with puromycin and after one week of selection, the cells expanded and cultured in triple flask and induced by doxycyclin. For each construct, 500 ml cell supernatants were collected and after filtration, the secreted Ebola proteins were purified by Strep-Tactin®XT 4Flow® column (IBA). After a high salt wash (1 M NaCl, 40 mM Tris-HCl, pH 8), the proteins were eluted (100 mM Tris-HCl, 150 mM NaCl, 1 mM EDTA, 50 mM biotin, pH 8) and then dialyzed against PBS. Protein concentration was corrected by the calculated extinction coefficients for the different proteins. The final viral proteins were stored at −80 °C.

### Protein G-based antibody purification

Transfected HEK293-6E cell cultures were harvested by centrifugation and cell-free supernatants were filtered followed by incubation with Protein G Sepharose beads (GE Life Sciences) at 4 °C overnight. Protein G Sepharose beads were transferred to columns and washed three times with PBS followed by elution of bound antibodies with 0.1 M glycine (pH 3). The resulting eluate was neutralized with 1 M Tris-HCl (pH 8). Buffer was exchanged to PBS with 30-kDa Amicon spin membranes (Millipore) and antibodies were quantified by UV/Vis spectroscopy (Nanodrop, A280). All antibodies were sterile filtered (0.22 μm, Millipore) and stored at 4 °C.

### Purification of serum IgGs

Serum was diluted 1:1 in Dulbecco’s PBS (DPBS) (Gibco, Thermo Fisher), and IgG was purified by three sequential passes over a 1.5 ml protein G agarose column (Pierce, Thermo Fisher). Following extensive washing with DPBS, bound IgG was eluted using glycine-HCl (pH 2.7) and immediately neutralized with 1 M Tris-HCl (pH 8.0). Buffer exchange to PBS and concentration of the purified IgG were carried out using 10-kDa molecular-weight-cutoff filters (Thermo Fisher).

### Determination of antibody concentrations by human IgG capture ELISA

Antibody concentrations in unpurified supernatants from transfected HEK293T or HEK293-6E cells were quantified using a human IgG capture ELISA, performed as previously described^31^ with the following minor modifications: ELISA plates (Greiner Bio-One) were coated with polyclonal goat anti-human IgG (2.5 µg/ml in PBS) overnight at 4 °C, followed by blocking with 2% bovine serum albumin and 0.1% Tween-20 in PBS for 60 min at room temperature. Cell culture supernatants (starting dilution 1:20) or serially diluted human myeloma IgG1κ standards (Sigma-Aldrich, I5154; 4 µg/ml) were then added. Bound antibodies were detected using horseradish peroxidase (HRP)-conjugated anti-human IgG (Jackson ImmunoResearch, 109-035-098, RRID: AB_237586; 1:2,500 dilution in blocking buffer). The reaction was developed with ABTS substrate (Thermo Fisher, 002024), and absorbance was measured at 415/695 nm (Tecan). Antibody concentrations were determined relative to the IgG1 standard curve.

### Cloning and production of pseudotyped lentiviral particles

Full filovirus GP coding regions (GeneBank ID: EBOV AF086833.2, BDBV FJ217161, SUDV FJ968794.1) with original leader peptides were codon optimized (Vectorbuilder Codon Optimization Tool) and ordered as gene fragments cloned into expression plasmids (pTwist CMV BG WPRE Neo) from Twist Bioscience. Lyophilized plasmids were solved in nuclease-free water (Ambion) at a final concentration of 100 ng/µl and pre-amplified by bacterial transformation. Pseudo-typed lentivirus was produced as described previously with minor modifications^38,39^. In brief, individual plasmids encoding HIV-1 Tat, HIV-1 Gag/Pol, HIV-1 Rev, Firefly luciferase followed by an IRES-ZsGreen, and the corresponding filovirus protein construct were co-transfected in HEK293-T cells 2-4 h after a change to FreeStyle 293 Expression Medium (Thermo Fisher) using FuGENE 6 Transfection Reagent (Promega) and incubated at 37 °C and 5% CO_2_. Culture supernatant was harvested at 40 h and 72 h post transfection, filtered by a 0.45 µM PVDF syringe filter and stored at −80 °C.

Virus titers were determined by infecting HEK293T cells with serial dilutions of viral supernatant. After 48 h of incubation at 37 °C and 5% CO₂, cells were lysed for 2 min with luciferin/lysis buffer (10 mM MgCl₂, 0.3 mM ATP, 0.5 mM Coenzyme A, 17 mM IGEPAL; all Sigma-Aldrich) containing 1 mM D-luciferin (GoldBio) in Tris-HCl. Luciferase activity was quantified using a microplate reader (Tristar 5, Berthold Technologies; 1 s counting time). For neutralization assays, viruses were diluted in FreeStyle medium containing 1% bovine serum albumin to yield relative light units (RLUs) approximately 1,000-fold higher than those of uninfected controls, corresponding to ∼50,000-100,000 RLUs.

### Pseudovirus neutralization tests

Neutralization activity of mAbs was first tested in a microneutralization assay in a BSL-2 facility. Serial dilutions of mAbs were prepared in DMEM (Gibco) containing 1 mM L-Glutamine in a 96-well format and incubated with pseudovirus for 1 h at 37 °C and 5% CO_2_. 1.25×10^4^ HEK293T cells were added per well and incubated at 37 °C and 5% CO_2_ for another 48 h. Luciferase activity was measured as described above. Negative controls (cells only and virus only) and virus positive controls (infected cells without mAb) were measured on each assay plate. The mean RLU of negative controls was subtracted as background signal and 50% inhibitory concentration (IC_50_) values were calculated as the mAb concentration leading to a 50% reduction of RLUs in comparison to the virus positive control.

### ELISA for antibody binding activity

High-binding ELISA plates (Greiner Bio-One) were coated with GP of EBOV, BDBV or SUDV (3 µg/ml), blocked with 2% bovine serum albumin/0.05% Tween-20 in PBS for 1 h at 37 °C and incubated with serial 1:4 dilutions of mAbs starting at 30 µg/ml. For detection of bound antibodies anti-human IgG-HRP (Jackson ImmunoResearch, 109-035-098, RRID: AB_237586; 1:2,000 in 2% bovine serum albumin/PBS) was incubated for 1 h at room temperature. Plates were washed with PBS/0.05% Tween-20 and developed with ABTS (Thermo Fisher, 002024), and absorbance was measured at 415/695 nm (Tecan). All samples were tested in duplicate.

### Antibody-competition ELISA

Selected mAbs were biotinylated using an EZ-Link Sulfo-NHS-Biotin kit (Thermo Fisher) and buffer exchanged into PBS with Amicon 10-kDa filters (Millipore). High-binding ELISA plates (Greiner Bio-One) were coated with EBOV-GP (3 µg/ml) overnight at 4 °C, blocked with 2.5% bovine serum albumin/2.5% nonfat milk/PBS for 1 h at 37 °C. Competing antibodies were added at 30 µg/ml and serially diluted 1:3, followed by the addition of 0.5 µg/ml biotinylated test antibodies (1 h at room temperature). Detection was performed with peroxidase-streptavidin (Jackson ImmunoResearch; 1:5,000 in PBS/0.5% bovine serum albumin/0.5% nonfat milk/0.05% Tween-20). Plates were washed with PBS/0.05% Tween-20 between steps, developed with ABTS (Thermo Fisher, 002024) and read at 415/695 nm (Tecan).

### Autoreactivity evaluations in HEp-2 cell assays

HEp-2 cell autoreactivity was assessed with a Kallestad HEp-2 kit (BIORAD) following manufacturer’s protocol. mAbs were applied and images were acquired using a Leica DMi6000B.

### Surface Plasmon Resonance (SPR) measurements

The binding of B10 and E9 Fabs to EBOV-GP was measured using a Biacore 8K instrument (Cytiva). EBOV ectodomain was first immobilized on a CM5 sensor chip (Cytiva) at a coupling density of ∼1,000 response units (RU). EBOV was immobilized on one of the two flow cells on each channel, where a non-relevant protein was immobilized on the second flow cell to serve as a blank. A single-cycle kinetics binding analysis was performed by injecting the Fabs at a series of concentrations (i.e., 20, 100, 500, 2,500, and 5,000 nM) in TBS-Tween buffer pH 8 (0.005% v/v Tween20) at a flow rate of 30 μL/min for a contact time of 120 s and final dissociation of 900 s. The sensor chip was regenerated using 4 mM NaOH.

### Cryo-EM image acquisition, data processing, and model building

A 3.5 μL sample of purified EBOV-GP at 0.4 mg/mL was mixed with 0.5 μL of B10 or E9 Fabs (at a 1:3 molar ratio) and kept on ice for 5 minutes. Samples were applied on a glow-discharged (8 s, 12 mA; Pelco easiGlow, Ted Pella) graphene oxide Quantifoil copper grids, R1.2/1.3, (Electron Microscopy Sciences) using a Vitrobot system (Thermo Fischer/FEI) (3.0 s blotting time, 4 °C, 100% humidity). Samples were incubated on the grid for 1 min before blotting was carried out. Cryo-EM data were collected on the Titan Krios microscope (FEI) operated at 300 kV, using a Gatan K3 direct detection camera. The beam size was 705 nm diameter (fringeless illumination), the exposure rate was 18 e s−1 pixel−1, and movies were then obtained at 105,000× magnification with a pixel size of 0.824 Å. The nominal defocus range was −0.8 to −2.0 μm. Data processing was carried out with the cryoSPARC v4.1.2 suite^40^. Patch motion correction and patch CTF estimation were carried out using cryoSPARC Live. Particles were extracted using a 256-pixel box size, and the data sets were cleaned and classified as illustrated for each map.

### Model building, refinement, and analysis

The initial model was generated using ModelAngelo^41^. This initial model was then manually completed and refined using Coot^42^ and real-space refinement in Phenix^43^. Structural analysis and representation were done using PyMol^44^, and ChimeraX^45^.

### Determination of antibody PKs *in vivo*

The half-lives of wild-type antibodies (B10, 10-1074, and 3BNC117) were assessed in NOD.Cg-*Rag1^tm1mom^Il2rg^tm1Wjl^*/SzJ mice (NRG; Jackson Laboratory; n = 12) following intravenous administration of 0.5 mg purified antibody in PBS. Total human IgG serum concentrations were monitored over 21 days by ELISA as described previously minor modifications^46^. Briefly, high-binding plates (Corning) were coated with anti-human IgG (2.5 μg/ml; Jackson ImmunoResearch), blocked with PBS containing 2% bovine serum albumin, 1 μM EDTA, and 0.1% Tween-20, and incubated with serial dilutions of a human IgG1κ standard (Sigma-Aldrich) or serum samples. Bound IgG was detected using HRP-conjugated anti-human IgG (1:2,000), developed with ABTS substrate, and absorbance was measured at 415 nm (Tecan). Plates were washed with PBS containing 0.05% Tween-20 between steps. Pre-dose samples confirmed the absence of human IgG prior to antibody administration. The mouse experiments were approved by the State Agency for Nature, Environment and Consumer Protection (LANUV) of North Rhine-Westphalia.

### Bacterial Strains

Escherichia coli DH5α strain (Thermo Fisher Scientific) was used for plasmid amplification of previously cloned expression vectors of orthoebolavirus-GP-targeting mAbs^13^.

### EBOV and SUDV neutralization assay

Antibodies were serially diluted in 96-well culture plates in DMEM supplemented with 2% fetal calf serum (FCS), penicillin (50 units/mL), streptomycin (50 µg/mL) (P/S) and glutamine (2 mM) (Q). A tissue culture infectious dose (TCID_50_) of 100 units of either SUDV (Boniface, GenBank: FJ968794) or EBOV (Mayinga, GenBank: NC_002549) was added to the serum dilutions in an equal volume of DMEM 2% FCS, P/S and Q. After incubation at 37 °C for 1 hour, approximately 10,000 Vero C1008 cells (ATCC CRL-1586) were added to each well. Vero C1008 cells were cultured as described elsewhere and authenticated in 2016 by DNA profiling of eight highly polymorphic regions of short tandem repeats by the “Leibniz-Institut DSMZ (Deutsche Sammlung von Mikroorganismen und Zellkulturen) GmbH”. They are proven to be free of mycoplasma through regular testing. Plates were then incubated at 37 °C with 5% CO_2_, and cytopathic effects (CPE) were evaluated at day 7 post infection. Neutralization was defined as complete reduction of CPE in antibody dilutions compared to positive controls. Neutralization titers of three replicates were calculated as geometric means (reciprocal value). The lower detection limit of the assay is determined by the first dilution of the respective antibody including the added virus. Neutralization assays were performed in the BSL-4 laboratory of the Institute of Virology, Philipps-University Marburg, Germany.

### Animal experiments

Male and female C57BL/6J IFNAR^−/−^ mice (9-24 weeks old), which lack the interferon-α/β receptor and are therefore susceptible to filovirus infection, were used as the animal model^28^. Animals were obtained from U. Kalinke (Twincore, Hanover, Germany) and subsequently bred in-house. All procedures were approved by the local authorities (animal welfare committee; Marburg: Regierungspräsidium Gießen, AZ V54-19 c 20 15 h 01 MR 20/7 Nr. G 64/2024) and conducted in accordance with Federation of European Laboratory Animal Science Associations (FELASA) and Gesellschaft für Versuchstierkunde / Society of Laboratory Animal Science (GV-SOLAS) guidelines, the German Animal Welfare Act, and Directive 2010/63/EU. Two weeks before the experiment, mice were allocated to experimental groups (2-3 animals per sex) and housed in isocages (Tecniplast) under SPF conditions following FELASA guidelines. At the same time they were tagged under brief isoflurane anesthesia (CP-Pharma) with an ear mark and implanted subcutaneously with a transponder (THERMOCHIP MINI, MSD) for body temperature measurements as described elsewhere^47^. On day −1, mice received an intraperitoneal injection of 40 mg/kg DZIF10C (placebo), B10, or mAb114, as previously described^48^. On day 0, mice were infected intraperitoneally with 1,000 PFU of EBOV or SUDV under brief isoflurane anesthesia. In addition to standard diet, all animals were provided access to a high-calorie energy gel after infection. Mice were monitored daily for body weight, body temperature, general condition, and spontaneous behavior, with each category scored with up to 10 points (total clinical score range: 0-40). Animals were euthanized under isoflurane anesthesia by cervical dislocation upon reaching humane endpoint criteria at a clinical score ≥10 or ≥6 on two consecutive days. Blood samples were collected on days 3, 7, 11 and 14 or at the humane endpoint, by facial vein puncture under brief isoflurane anesthesia (Microvette® CB 300 Serum CAT, Sarstedt).

### Quantitative real-time RT-PCR analysis of virus load in mouse tissue samples

Tissue and serum samples were processed as described elsewhere. RT-qPCR was performed using the RealStar® Filovirus Screen RT-PCR Kit 1.0 (Altona Diagnostics, Cat. No. 441013) following the manufacturer’s instructions, and reactions were run on a qTOWER³ instrument (Analytik Jena).

### Virus titration by TCID_50_

Tissue and serum samples were processed as described elsewhere. To quantify infectious virus in organ and serum samples, 10⁴ Vero C1008 cells/well were seeded into 96-well plates. On the following day, cells were inoculated with 5-fold serial dilutions of supernatants from infected cells or organ homogenates. 7 days post infection, CPE were assessed microscopically, and TCID₅₀/ml values were calculated according to the method of Spearman-Kärber.

### Quantification and statistical analysis

Flow cytometry data were analyzed and quantified using FlowJo (v10). Quantitative and statistical analyses were carried out in GraphPad Prism (v10), Microsoft Excel for Mac (v16.99), and Python (v3.6.8). Sequence alignments were performed using IgBlast and Clustal Omega (v1.2.3).

### Use of large language models

ChatGPT (v.4 and v.5) was used for general editorial tasks, including proof-reading, grammar correction and text summarization only. Scientific content and conclusions are the work of the authors.

